# The Hox gene *Antennapedia* regulates wing development through 20-hydroxyecdysone in insect

**DOI:** 10.1101/2021.05.26.445904

**Authors:** Chunyan Fang, Yaqun Xin, Tao Sun, Antónia Monteiro, Zhanfeng Ye, Fangyin Dai, Cheng Lu, Xiaoling Tong

## Abstract

A long-standing view in the field of evo-devo is that insect forewings develop without any Hox gene input. The Hox gene *Antennapedia* (*Antp*), despite being expressed in the thoracic segments of insects, has no effect on wing development. This view has been obtained from studies in two main model species, *Drosophila* and *Tribolium*. Here, we show that partial loss of function of *Antp* resulted in reduced and malformed adult wings in *Bombyx, Drosophila*, and *Tribolium*. *Antp* mediates wing growth in *Bombyx* by directly regulating the ecdysteriod biosynthesis enzyme gene (*shade*) in the wing tissue, which leads to local production of the growth hormone 20E. In turn, 20E signaling also up-regulates *Antp*. Additional targets of *Antp* are wing cuticular protein genes *CPG24, CPH28*, and *CPG9*, essential for wing development. We propose, thus, that insect wing development occurs in an *Antp*-dependent manner.

## Introduction

The Hox genes encode a family of transcriptional regulators that are important in differentiating the bodies of bilaterian animals along their antero-posterior axis [1]. Disruptions to individual Hox genes often leads to disruptions of traits that develop in the regions where the Hox gene is expressed [1].

In holometabolous insects, the Hox gene *Antennapedia* (*Antp*) is expressed in all thoracic segments, including in the forewing and hindwing yet, no function has been attributed to this gene regarding wing morphogenesis. In *Drosophila*, wing and haltere primordia could be detected in embryos even in the complete absence of *Antp* function in homozygous mutants of *Antp* [2]. In addition, a very low level of Antp protein present in the growing wing imaginal disc, suggested that forewing formation did not require *Antp* [2]. Similarly, no obvious phenotypes were observed in adult *Tribolium* elytra (forewing) or hindwing after RNA interference (RNAi) of *Antp*. These data suggested that wing development takes place without any *Antp* input [3]. In contrast, the Hox gene *Ultrabithorax (Ubx)*, expressed exclusively in hindwings, functions to differentiate hindwings from forewings. Forewing development in insects was, thus, thought to occur without any significant Hox gene input [3–13].

Recently, however, we observed that two loss of function mutations in the silkworm *Bombyx mori Antp* gene (*BmAntp*), *Nc* and *Wes*, displayed abnormal wings [14,15]. These mutations had not been examined beyond the embryonic stage due to lethality, but could be maintained in heterozygous lines. The adults of these lines displayed reduced and malformed wings.

These two *Bombyx Antp* mutants, however, shared common features with *Drosophila Antp* mutants, observable in embryos. The homozygous (*Antp*^-/-^) embryos died late in embryogenesis but displayed a homeotic transformation of thoracic legs to antenna-like appendages[14–17]. The novel wing phenotypes in *Bombyx* heterozygote mutants, however, suggested that *Antp* was affecting wing development, a role not previously documented for this gene because embryos died long before the stage when wings start to develop. In the present study, we used wiltype and heterozygotes of the *Wes* strain (*Antp*^+/−^) as the study objects to more fully undertstand the role of *Antp* in wing development.

## Results

### *BmAntp* is involved in the development of wings in *Bombyx*

Since defective adult wings were observed in aberrant *Antp Wes* and *Nc* mutants (*Antp*^+/−^) [14,15], we sought to test when in development *Antp* input was required. We analyzed the expression profile of *BmAntp* in the forewing and hindwing of wildtype (WT) individuals from the 3^rd^ day of the 5th instar to the adult stage. qRT-PCR revealed that the expression of *BmAntp* was maintained at a low level in the larval stage and gradually increased and reached a peak on the 6^th^ day of the pupal stage. Forewings expressed higher levels of *Antp* relative to hindwings at most times during the pupal stage (Fig 1A). Then, we compared the expression pattern of *BmAntp* between mutant *Wes* (*Antp*^+/−^) and WT individuals. *BmAntp* was expressed at a consistent but lower level in the mutants compared to WT controls (Fig 1B).

**Fig 1.**
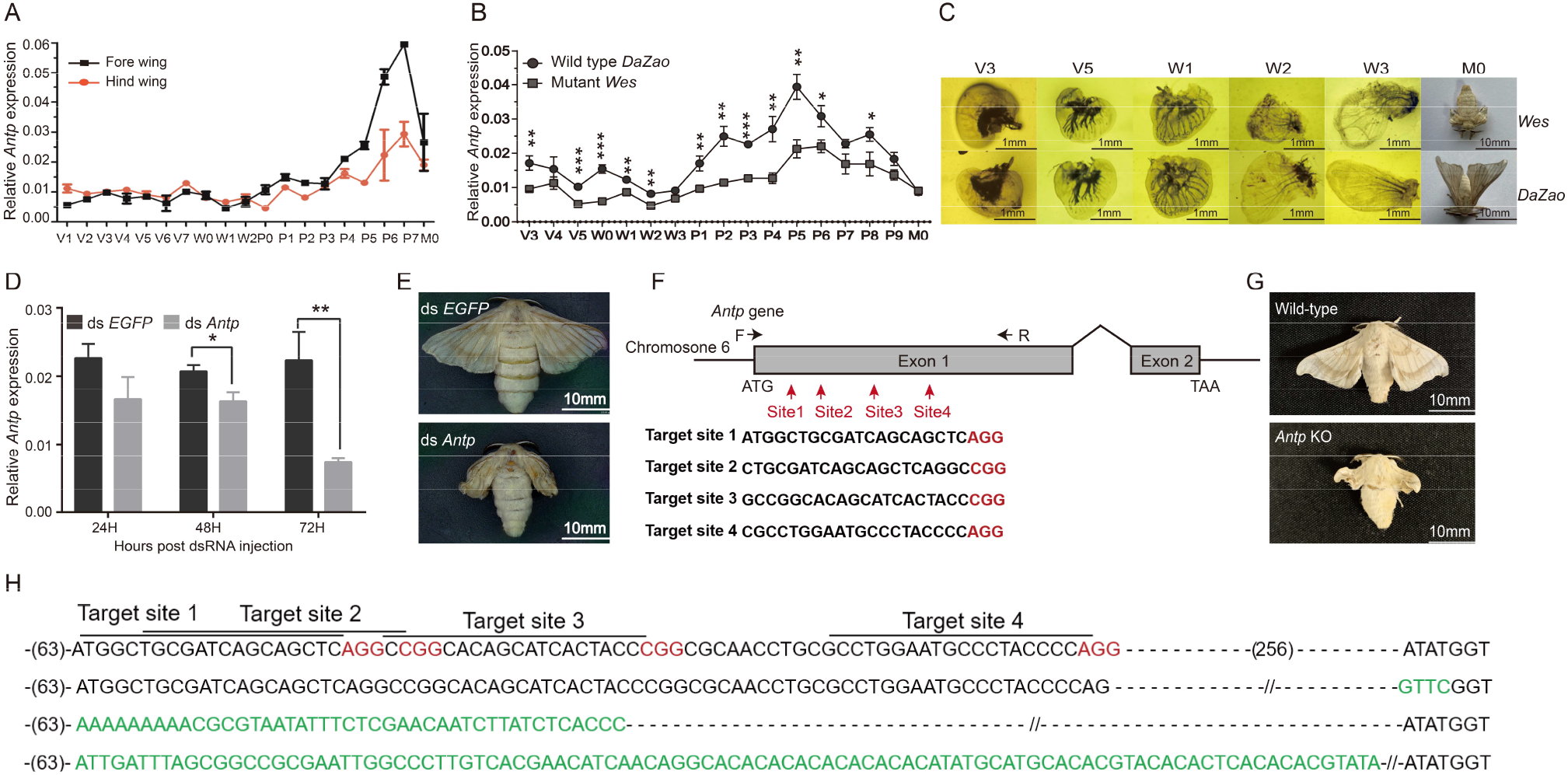
*Antp* is essential for wing development in *B. mori*. (A) Temporal expression pattern of *Antp* in wild-type (*DaZao*) forewing and hindwing discs by qRT-PCR. (B) Expression profiles of *Antp* in the wing discs of wild-type and mutant (Wes) lines from larvae to adult stages. (C) Phenotype of the wing discs in wild-type *DaZao* and *Wes* (*Antp*^+/−^) mutants over different time points. (D) Relative *Antp* expression levels of dsRNA-treated larvae at 24 h, 48 h, and 72 h after dsRNA treatments. Animals injected with *dsEGFP* served as controls. (E) Wing phenotype of dsRNA-treated silkworm adults. (F) Genomic structure of *Antp*. The single guide RNA (sgRNA) target sequence is in black font and the protospacer adjacent motif (PAM) sequence is in red font. The red arrows mark the sgRNA targets on the *Antp* gene. F and R indicate the approximate locations of the amplification primers. (G) Representative phenotypes of wild-type (top) and mutated (bottom) insects, with smaller and abnormal wings. (H) Mutated sequences of crispant individuals. The wild-type sequence, showed above the mutant sequences, is in black and the PAM sequence is in red font. The size of indels is shown to the right of the sequence. Inserted sequenced are in green font. For all graphs, V is the 5th instar larvae, V1–V7 means days 1–7 of the 5th instar larvae; W is the wandering larval stage, W0–W3 indicates days 0–3 of the wandering larval stage; P, the pupal stage, P0–P9 indicates days of 0–9 of the pupal stage; M0, newly emerged adult. All experimental data shown are means ± SE (n=3). Asterisks indicate significant differences with a two-tailed t-test: *P<0.05, **P<0.01, ***P<0.001.

To evaluate the effects of *BmAntp* expression levels on wing morphology during development, we dissected the wing discs of *Wes* (*Antp*^+/−^) and WT from the 3^rd^ day of the last larval instar to the wandering stage larva. Wing disc size increased slowly during the larval stage and was not significantly different between *Wes* mutants (*Antp*^+/−^) and WT individuals. Then, during the wandering stage, the wing morphology changed dramatically. The wing discs of *Wes* mutants (*Antp*^+/−^) were curlier and smaller than those of WT, and finally degenerated to tiny and wrinkled adult wings (Fig 1C).

To confirm the function of *BmAntp* in wing development, we performed RNAi injections into WT larvae of *B. mori*. We synthesized dsRNA targeting *BmAntp* and injected it into larvae on the 1^st^ day of the wandering stage. qRT-PCR showed that *BmAntp* dsRNA efficiently reduced *BmAntp* transcript levels compared to controls injected with *EGFP* dsRNA (Fig 1D). Nineteen out of 22 (86%) *BmAntp* dsRNA treated individuals had small wings similar to the *Wes* mutant (*Antp*^+/−^) while control silkworm adults grew their wings normally (Fig 1E).

To further confirm the function of *BmAntp* we performed crisper-Cas9 injections into WT embryos of *B. mori*. We generated a genomic disruption of the *BmAntp* gene by targeting its first exon using four specific single-guide RNAs (sgRNAs) and the Cas9/gRNA ribonucleoprotein (RNP) delivery system (Fig 1F). After injection, 20 eggs hatched and 18 larvae developed to the adult stage. We found that 61% of the moths (11 individuals) displayed malformed adult wings (S2 Table, Fig 1G), and confirmed that various insertions and deletions were present at the location targeted by the four sgRNAs (Fig 1H). Abnormal wings were not observed in control injections with *BmBLOS2* sgRNA which only led to translucent larval skin. These data indicate that *BmAntp* is a critical transcription factor that regulates wing development in *B. mori*.

### BmAntp affects the synthesis of 20E by regulating the expression of *Shade* in wing discs

We next tested whether the production of abnormal wings in *BmAntp* mutants was related to deficits in levels of the molting hormone, 20-hydroxyecdysone (20E). We tested this hypothesis because 1) Significant differences in the size of wing discs were observed between *BmAntp* mutants and controls starting from the onset of the larva-to-pupa transition (Fig 1C); 2) A pulse of this steroid hormone normally regulates the larva-to-pupa transition; and 3) 20E is a major regulator of wing growth and development [16, 17].

We first examined the expression level of genes involved in the ecdysteriod biosynthesis pathway in a variety of tissues. We found that *spookier, phantom, disembodied*, and *shadow* were expressed in the prothoracic gland (PG), as expected, as this is the main source of ecdysteroid synthesis in insect larvae [5].In addition, the *shade* gene, which codes for a P450 monooxygenase that catalyze ecdysone into the active 20E in targeted peripheral tissues [18], was primarily expressed in the wing discs compared to the PG and hemolymph (Figs 2A–2E).

**Fig 2.**
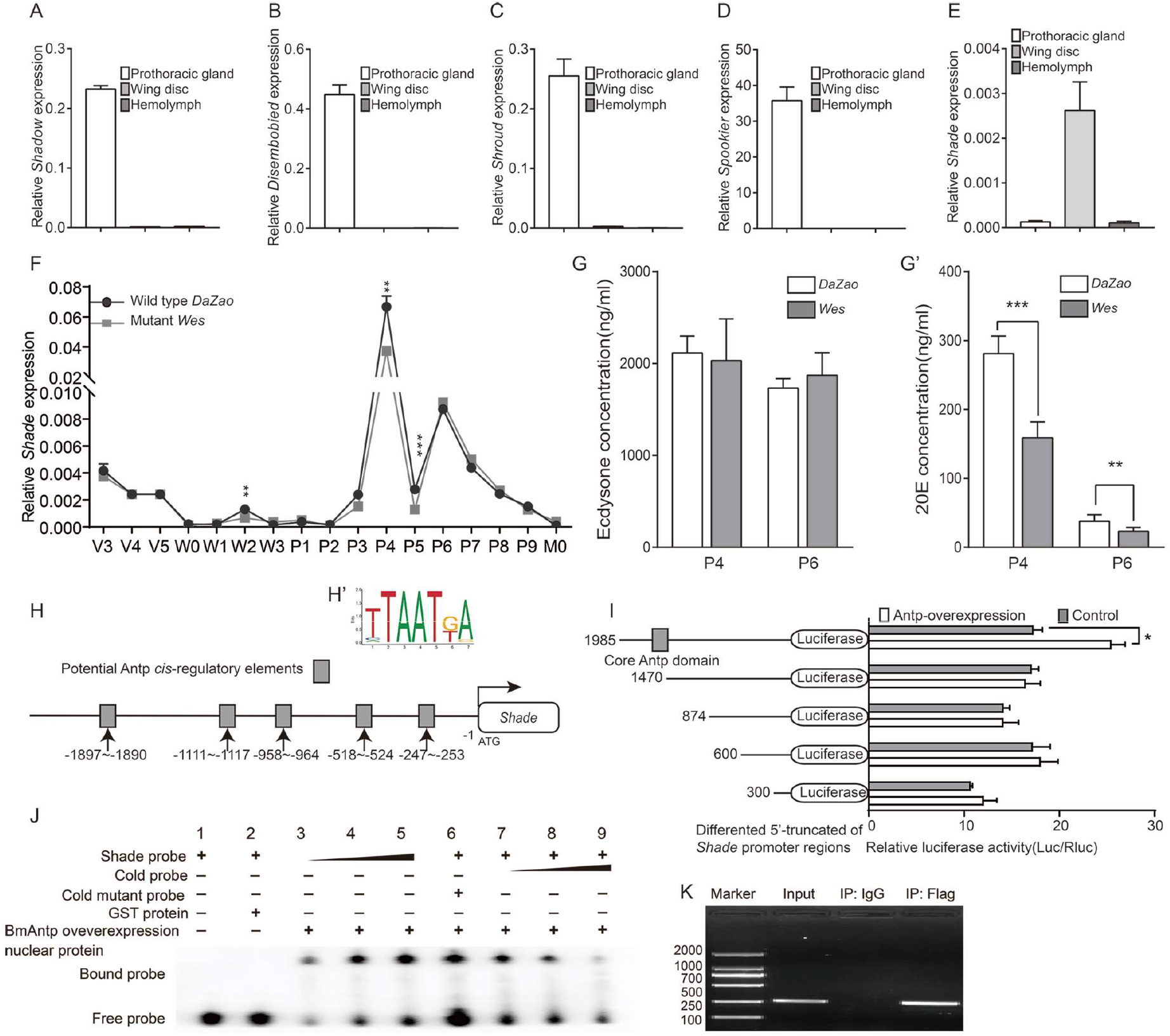
*Antp* induces 20E synthesis in the wing tissue by directly binding to the *shade* promoter. (A–E) Relative expression of five ecdysteriod enzyme genes in the prothoracic gland, hemolymph, and wing disc. (F) mRNA levels of *shade* were detected by qRT-PCR from the 5th instar larval stage to the adult stage. (G and G’) The titers of ecdysone (G) and 20E (G’) in *Bombyx* wing discs of WT *DaZao* and *Wes* mutants (*Antp*^+/−^) at P4 and P6. (H) Location of the five potential Antp binding sites in the *shade* promoter. (H’) Classic Antp binding motif. (I) The effect of different truncations of the *shade* promoter on luciferase activity when Antp is overexpressed in BmN cells. (J) EMSA confirmed that the recombined Antp proteins bind to the nt −1897–-1890 region in the *Shade* promoter. Coincubating nucleoproteins from *Escherichia coli* strain BL21 (DE3) competent cells overexpressing GST with labeled Antp probes resulted in loss of the binding band. Purified recombinant BmAntp protein could bind to the biotinylated probes in a dose-dependent manner (lanes 3–5), and this binding could be competitively suppressed by unlabeled probe (lane 6). The unlabeled probe with mutation in the core-binding motif of BmAntp could not compete for BmAntp binding to biotinylated probes (lanes 7–9). We further validated the direct regulation of BmAntp on shade transcription through in vivo ChIP-PCR following the BmN cells which were overexpression of FLAG-tagged BmAntp. (K) ChIP-PCR assay of the direct binding of Antp to the *shade* promoter in BmN cells with Antp-Flag overexpression. Specific primers covering Antp binding sites of the 3 *Shade* promoter were used. Comparing with nonspecific IgG antibody, used as a negative control, the antibody against FLAG can specifically immunoprecipitate the DNA regions including −1985 to −1470 of the Shade promoter. All experimental data shown are means ±SE (n=3). Asterisks indicate significant differences with a two-tailed t-test: *P<0.05, **P<0.01, and ***P<0.001.

We next explored whether *Wes* mutants (*Antp*^+/−^) expressed *shade* at different levels relative to WT wings, and whether this impacted levels of 20E in the wing tissue. The *shade* transcripts were present at markedly higher levels in WT than in *Wes* mutant (*Antp*^+/−^) wings, and levels reached a peak on the 4^th^ day of the pupal stage (Fig 2F). Titers of ecdysone measured from wing discs on that day (P4), were similar between *Wes* mutants (*Antp*^+/−^) and WT individuals. Titers of 20E, however, were significantly lower in *Wes* mutants (*Antp*^+/−^) relative to WT individuals two days later (on P6) (Figs 2G and 2G’).

We next investigated whether the expression levels of *Ecdysone Receptor (EcR*) and *ultraspiracle (usp*) [19], the receptors that bind 20E to transduce edysone signaling to the nucleus were also different between *Wes* and WT individuals. This is because 20E signaling is known to up-regulate expression of *EcR* and *usp* in the wings of *Drosophila* [20,21]. Significantly lower levels of *usp*, and of the two isoforms of *EcR, EcRA* and *EcRB* mRNA were detected in the mutants compared with WT on day P4 (S1 Fig). These results suggest that Antp is also regulating the expression of these genes, either directly or indirectly. The latter mechanism could involve Antp up-regulating *shade*, which increases 20E titers in the wing cells which, in turn, up-regulates *EcR* and *usp* transcription in wings.

We next sought to test whether *shade* was a direct target of BmAntp. We examined a 2 kb region of DNA immediately 5’ of the start site of *shade* for possible Antp binding domains and found a total of five such domains (Figs 2H and 2H’). To evaluate the extent that DNA containing one or more of these domains could regulate flanking gene expression we cloned different sized fragments, containing a different number of Antp binding domains, upstream of the reporter gene *luciferase*. We transfected this plasmid into *Bm*N cells and co-transfected *BmAntp* in these cells as well (S2 Fig). The largest fragment (−1985 to −300), containing all five Antp binding sites, led to significantly increased luciferase activity compared to the other four fragments (Fig 2I). These data suggest either that a regulatory region −1985 to −1470 containing a key Antp binding site or, more likely, that all Antp sites together are required for the transcriptional regulation of *shade*, and that *shade* is likely a direct target of BmAntp.

To determine whether BmAntp protein could directly bind to the *in silico* identified Antp binding sites of the *shade* promoter, we designed a specific biotinylated probe covering the −1985 to −1470 genomic region of *shade* and conducted electrophoretic mobility shift assay (EMSA) (Fig 2J). We further validated the direct regulation of BmAntp on Shade transcription through *in vivo* ChIP-PCR following the BmN cells which were overexpression of FLAG-tagged *BmAntp* (Fig 2K, S3 Fig). Our data indicated that BmAntp activates the transcription of *shade* by directly binding to the tested genomic region.

### *BmAntp* is upregulated by 20E

We next sought to investigate which genes could be driving *Antp* expression in the wings of *B.mori. Antp* levels were low throughout wing disc development until the pupal stage, and then followed a slow rise and fall. Because this expression profile resembled the 20E titer profile in *B.mori* hemolymph [22], we decided to investigate whether the 20E/EcR/USP complex could be upregulating *Antp* in pupal wings. We first examined potential Ecdysone Response Element (EcRE) binding sites for the complex within the ~2 kb upstream of *BmAntp* (counting from the start codon of 5’ UTR) and discovered three such sites (EcRE1, −139– −153 nt; EcRE2, −1034– −1048 nt; EcRE3, −1592– 1606 nt) (Fig 3A). Then, we cloned this ~2kb genomic region of *BmAntp* in front of the luciferase reporter gene and transfected this plasmid into BmN cells, followed by 20E treatment. Subsequent dual luciferase reporter assays revealed that this region drove significantly higher luciferase activity after 20E application (Fig 3B). To further confirm the upregulation of *BmAntp* expression by 20E, we used qRT-PCR to show that the expression of *BmAntp* was significantly upregulated both in cultured cells and in wing discs after 20E treatment (Figs 3C and 3D). These data show that 20E can upregulate expression of *BmAntp* via the direct binding of the 20E/EcR/USP complex to a 5’ upstream region of the gene.

**Fig 3.**
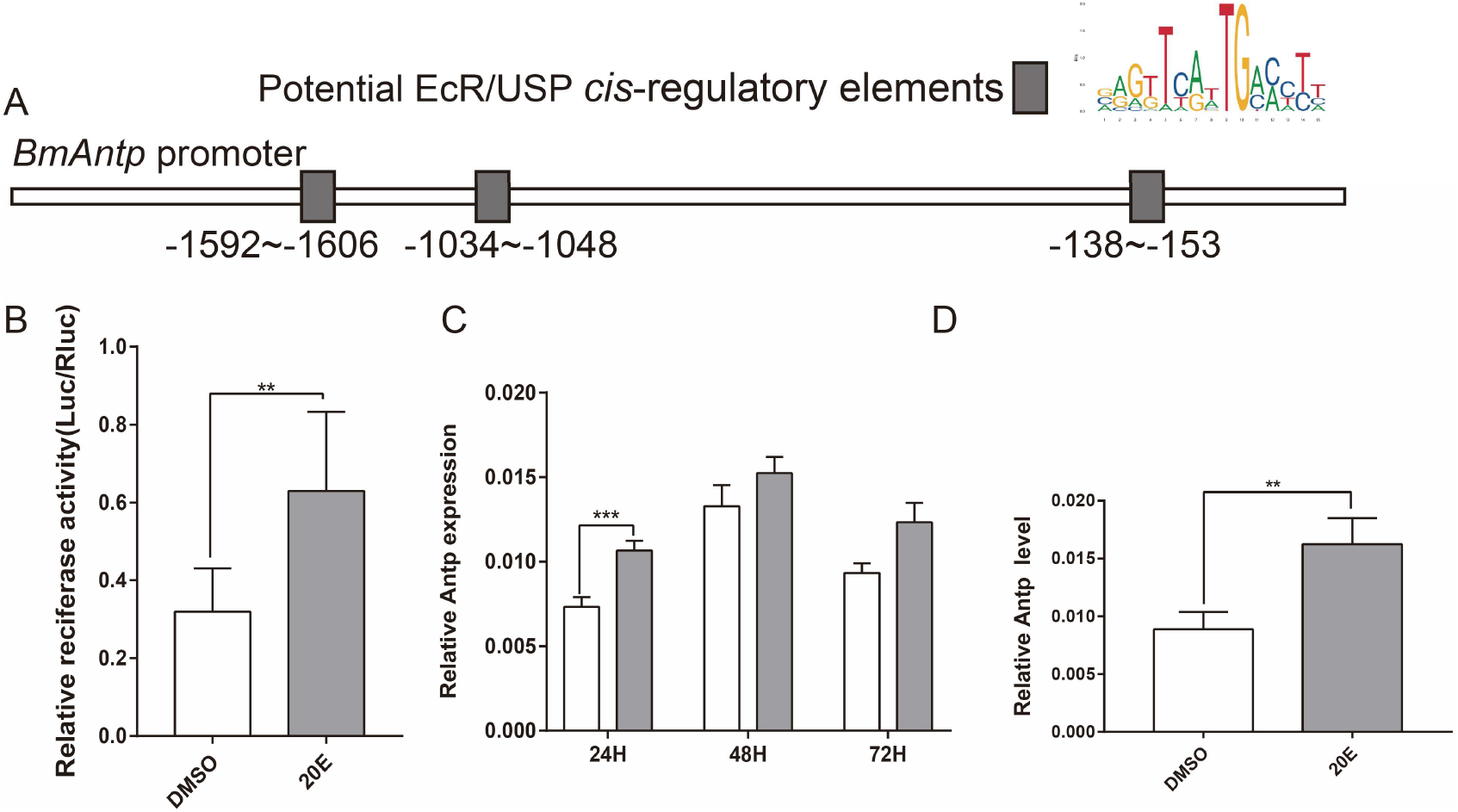
20E induced expression of Antp. (A) Location of potential Antp binding sites in the *Antp* promoter. (B) Effects of 20E treatment on the luciferase activity driven by the *Antp* promoter. (C and D) Levels of *Antp* expression increase in BmN cells (C) and wing discs (D) after 20E treatment. All of the experimental data shown are means ±SE (n=3). Asterisks indicate significant differences with a two-tailed t-test: **P<0.01.

### BmAntp directly regulates wing-specific cuticular protein genes

In order to explore potential additional targets of Antp, besides *shade*, that might have contributed to the small wings of adult *Wes* mutants, we investigated the expression of four cuticular proteins with a known expression profile, that matched that of *Antp*, in both WT and *Wes* mutants. In particular, expression levels of *CPH28, CPG24, CPG9* peaked at P5, as did expression of Antp (Fig 1B) [20]. *CPG11*, by contrast, was expressed primarily during the early 5^th^ instar, and was used as a control gene [23]. Previous work has shown that cuticular proteins are major components of insect wings and that both EcR-mediated signaling as well as other transcripton factors regulate their very dynamic and specific expression profiles [24–26]. qRT-PCR analysis showed that the expression levels of *CPH28, CPG24, CPG9*, and *CPG11* in *Wes* (*Antp*^+/−^) were remarkably lower than those of WT (Fig 4A). We explored the direct regulation of these four cuticular proteins by Antp by conducting Luciferase reporter assays in BmN cells with candidate genomic regions (3kb upstream of each gene) containing putative Antp binding sites (Fig 4B). Increasing *BmAntp* levels in these cells significantly upregulated the transcription of *CPH28, CPG24*, and *CPG9* (Figs 4C–4F), but not *CPG11*. A dual-Luciferase assay with *CPH28* further showed that BmAntp can directly elevate the expression of *CPH28* (Fig 4G). Moreover, an EMSA and ChIP-PCR essay showed that BmAntp was able to directly bind the *in silico* identified Antp binding sites in the *CPH28* promoter (Figs 4H and 4I, S4 Fig). These results indicate that BmAntp can upregulate the transcription of these three wing cuticular protein genes, and *CPH28* is likely up-regulatd by a direct interaction of Antp with this gene’s promoter.

**Fig 4.**
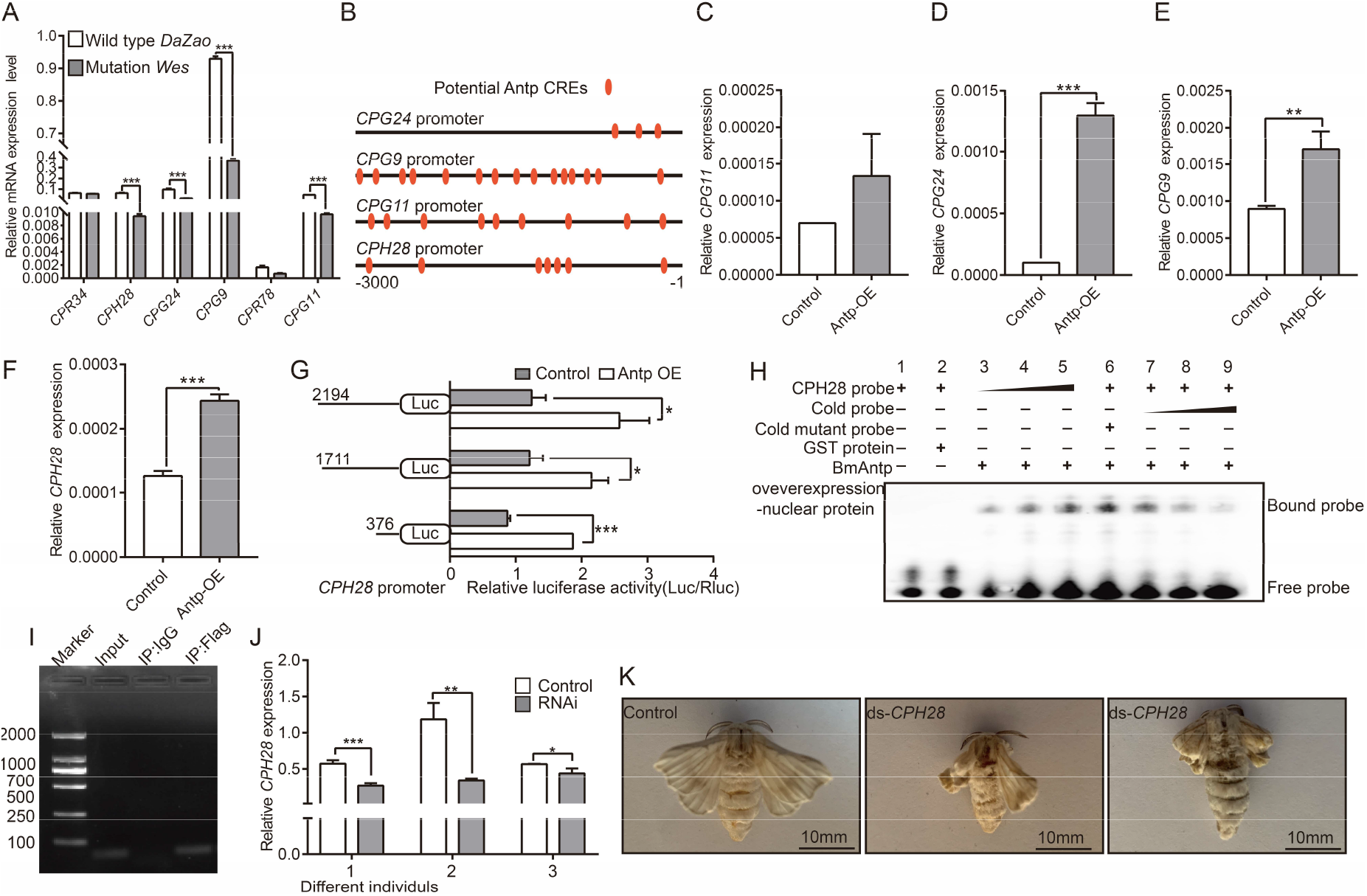
*Antp* regulates the expression of cuticular protein genes essential for wing development. (A) mRNA levels of cuticular protein genes in wing discs of *DaZao* and *Wes* (*Antp*^+/−^) at P5. (B) Schematic of the potential Antp CREs in the promoters of cuticular protein genes. (C–F) Relative cuticular protein genes expression detected in Antp overexpression BmN cells. (G) Antp increased luciferase activity driven by different truncations of the CPH28 promoter. (H) Electrophoretic mobility shift assay (EMSA) of the binding nuclear proteins extracted from Antp-overexpressing *Escherichia coli* strain BL21 (DE3) competent cells with the Antp binding motif. Co-incubating nucleoproteins from *E. coli* strain BL21 (DE3) competent cells overexpressing glutathione S-transferase (GST) with labeled Antp probes results in loss of the binding band. The binding signal between recombinant GST-BmAntp protein and Antp binding motif probe was gradually enhanced with increased probe levels (lanes 3–5). (I) ChIP-PCR assay shows that Antp binds directly to Antp binding motifs present in the *CPH28* promoter in BmN cells. A Flag tag was fused to BmAntp and an anti-Flag tag antibody was used in the ChIP assay. The cells were transfected with recombinant plasmid Flag-BmAntp, and then the cells were collected for ChIP assay 48 h post-transfection. The results showed that the anti-Flag antibodies, but not IgG (a negative control), precipitated DNA containing the Antp binding motifs in the cells transfected with the Flag-BmAntp expressing plasmid. (J) qPCR analyses of *CPH28* expression in wing discs of different individuals, 48 h after knock down of *CPH28* and of a control sequence (containing the scrambled siRNA sequence). (K) Comparisons of adult wing morphology after dsRNA injections. All experimental data shown are means ± SE (n=3). Asterisks indicate significant differences with a two-tailed t-test: *P<0.05, **P<0.01, and ***P<0.001.

To determine whether *CPH28* is essential for wing development, we knocked it down using RNAi. CPH28-siRNA was injected into 18 pupae, and the same quantity of scrambled siRNA sequence was injected in control animals. Levels of *CPH28* decreased significantly in the wing discs 48 h after *CPH28*-siRNA injections relative to control injections (Fig 4J). The ratio of malformed wings reached 80% after eclosion (Fig 4K, S3 Table). In contrast, all moths in the control group had normal wings (Fig 4K). These results indicate that *CPH28* is required for the generation of normal wings in silkworms.

### *Antp* function in wing development is conserved in *Drosophila* and *Tribolium*

To evaluate whether the function of *Antp* in wing development is conserved across other insect orders, we examined the wings of adult flies and beetles after *Antp* down-regulation. In *Drosophila* we drove expression of *Antp* RNAi hairpins in larval and pupal wing discs under the control of the *nubbin-gal4 (nub-gal4*) driver. All individuals in which *Antp* was knocked down had rudimentary wings that were reduced in size compred to controls (Figs 5A–5D). In *Tribolium*, we injected *Antp/ptl* dsRNA during the last larval stage, just before the onset of rapid wing growth [3]. These injections led to lower mRNA levels of *Antp/ptl* (S5A Fig) and to wrinkled and shortened forewings (elytra) and hindwings (Figs 5E–J, S5B and S5C Figs). Additionally, the uniform mesonotum phenotype observed in the *Antp*/*ptl* RNAi adults was consistent with that reported by Tomoyasu and colleagues (Figs 5K and 5L, S5D and S5E Figs) [3]. These observations indicate that *Antp* plays a crucial role in the development of wings in *Drosophila* and *Tribolium*. Taken together, these results demonstrate that *Antp* participates in insect wing development in a conserved manner.

**Fig 5.**
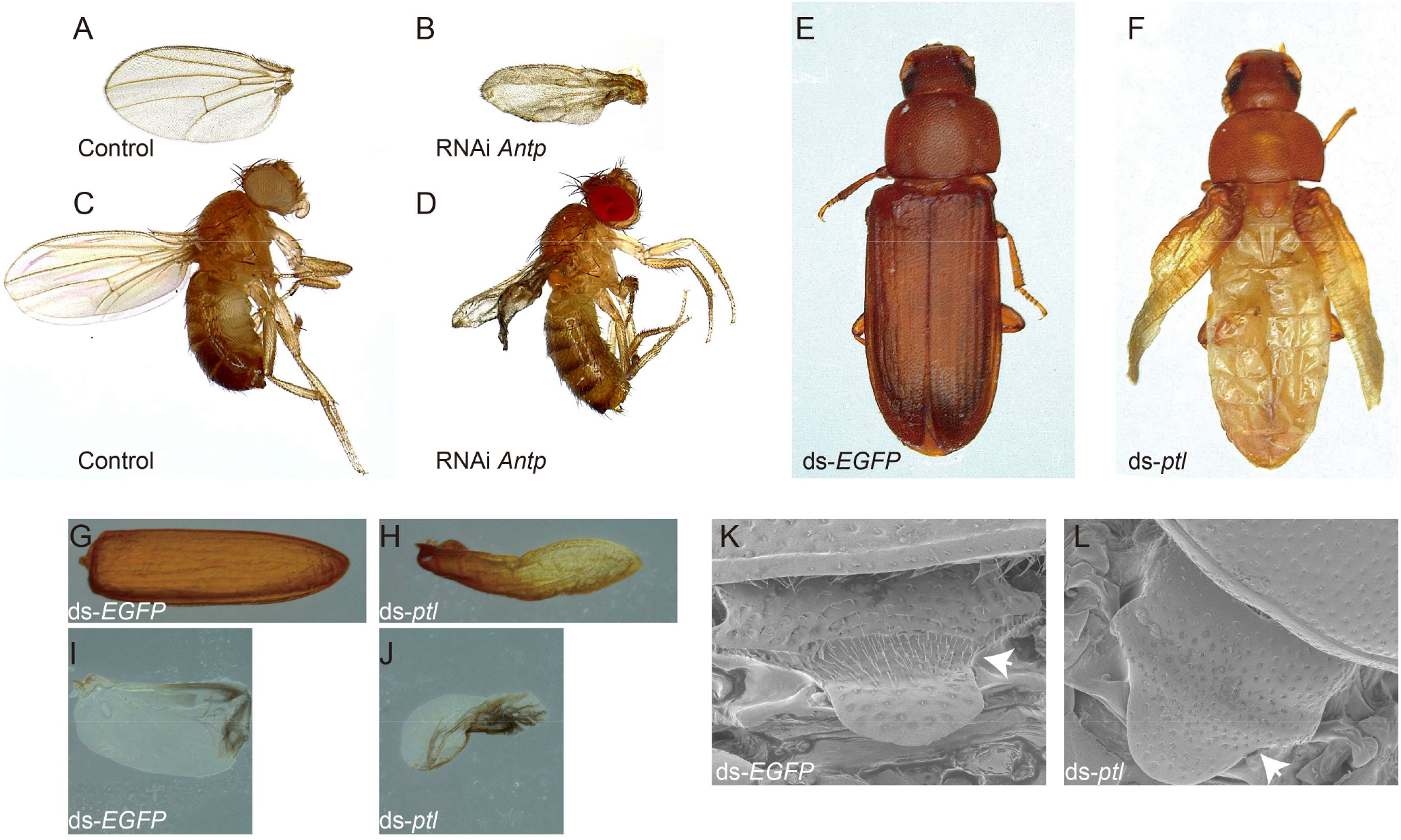
*Antp* is essential for wing development in *Drosophila* and *Tribolium*. (A and B) Adult *Drosophila* wings from control (A) and *Antp* RNAi (B) treated individuals. (C and D) Adult *Drosophila* of control (C) and *Antp* RNAi (D) treated individuals. (E and F) *ptl* RNAi leads to reduction of elytra and hindwings in *Tribolium* adults. (E) *ds-EGFP*. (F) *ds-ptl*. (G and I) The elytron (G) and hindwing (I) from a *ds-EGFP* treated individual. (H and J) The elytron (H) and hindwing (J) from a *ds-ptl* treated individual. (K and L) *ptl* RNAi leads to a uniform mesonotum (white arrow in K and *L)*. (K) *ds-EGFP*. (L) *ds-ptl*.

## Discussion

### Hox gene *Antp* is indispensable for wing development

Limited experiments in previous *Drosophila* studies, focusing on embryonic and larval stages, likely prevented the identification of Antp’s role in later stages of wing development. Fly embryos homozygous for *Antp*^W10^, a mutation in the *Antp* sequence, led to normal wing primordia, whereas ectopic expression of *Antp* in third instar larval wing discs had no effect on larval wing discs morphology [2]. This lack of results is expected as Antp protein was largely absent in the major region of the growing larval discs [2]. In the present study, the *nub-Gal4* driver was used to drive *UAS-Antp^RNAi^* expression in fly wing discs. We chose this driver as its expression was first detected in late 2nd instar wing discs and persisted through late pupal wings [27]. This led to a prolonged silencing of *Antp* expression and to malformed adult wings in *Drosophila*. Given that the silkworm *Antp* was also expressed at low levels in larval wings, but at much higher levels in pupal wings, we speculate that there is no requirement for *Antp* function during the embryo and larval stages, but *Antp* is important for wing development in the later pupal stages.

Our RNAi experiment in *Tribolium castaneum* also identified strong wing defects not previously identified with a previous similar RNAi experiment [3]. This previous study only reported variation of mesonotum morphology [3], which was also found in our experiment. We preformed the *Antp* RNAi experiment twice (>250 individuals) and obtained consistent defective wing morphologies, that were not observed in control animals injected with dsRNA against *EGFP*. We speculate that the different outcomes of the two experiments may be due to the different ds*Antp* fragments used. We used two fragments covering a larger region of the *Antp* gene (922bp) compared to the 535 bp fragment used by Tomoyasu et al.. Based on the present results, we propose that *Antp* is necessary for wing development in *Bombyx*, *Drosophila*, and *Tribolium*.

Recently, *Antp* input was found to be required for the development of two novel traits in the wings of the nymphalid butterfly *Bicyclus anynana:* silver scales and eyespot patterns, in both forewings and hindwings, but only minor wing growth deformities were reported (see Fig 2B in Matsuoka and Monteiro) [28]. It is possible that the role of *Antp* has shifted from a general wing growth role to a more specialized role in color pattern formation. This might be the case in this species and other nymphalids where *Antp* expression has been visualized in the eyespots [27,29]. Alternatively, the mosaic disruptions obtained with this crispr-Cas9 experiment were insufficient to uncover a more general role of *Antp* in wing growth and development. Most interestingly, the effect of Antp on *shade* expression shoud be investigated in connection to 20E-mediated eyespot size plasticity in this species [27,29].

### Bi-directional regulation between *Antp* and 20E

We showed that Antp directly binds to the promoters of *shade*, a gene coding for the last step in the production of the active ecdysteroid, 20E, and that 20E was produced inside wing tissues from the precursor ecdysone produced in the prothoracic gland [25,30]. The biosynthesis of 20E, the main hormonal regulator or molting and methamorphosis in insects [17,31], is mediated by the Halloween genes, such as *spookier, shroud, disembodied, shadow* and *shade* [32]. *shade* is known to converts ecdysone into 20-hydroxyecdysone (20E) in peripheral organs such as the fat body, midgut and Malpighian tubules [25,30]. As expected, the mRNA coding for *Shade* mRNA was present at an extremely low level in the prothoracic gland and also in the hemolymph, but at a higher level in wing discs. Given that the mRNA expression of *shade* in wing discs of *Wes* (*Antp*^+/−^) mutants was significantly lower than that in normal wing discs, this explains the observed lower levels of 20E, but not of ecdysone, in the wing tissue of these mutants, and associated wing disc growth disruptions.

In our study, we also found that supplementary 20E up-regulated *Antp* in both BmN cells and in developing wing discs. Similar regulation of Hox genes by 20E have previously been reported in the *Drosophila* heart, where the expression of *Ubx* and *abdominal-A (abdA)*, was also activated by ecdysone signaling [33]. So, Antp upregulates 20E in the wing, and 20E together with its nuclear co-receptors (EcR and USP) upregulate *Antp, EcR* and *usp* expression.

### Antp regulates the expression of wing cuticular protein genes

Cuticular proteins are major components of insect wings and previous studies had already implicated the regulation of these proteins by other Hox genes. A total of 52 cuticular protein genes were detected in silkworm wing discs by expressed sequence tags [23]. The regulation of one these proteins, BmWCP4, was previously shown to be dependent on the co-binding of the Hox gene *BmAbd-A* with the transcription factor *BmPOUM2*, in the gene’s promoter [34]. In the present study, we focused on investigating wing cuticular protein genes whose expression patterns were largely congruent with that of *Antp* [31]. We showed that they were remarkably down-regulated in mutant (*Antp*^+/−^) individuals, and that disruptions to one of these proteins impaired wing development. It remains possible, that many more additional *Antp* targets remain to be described.

Previous studies have assumed that the forewing is a Hox-free wing [3,5,9]. Our data indicated that *Antp* is crucial for wing development in insects (Fig 6). It does this by directly enhancing transcription of the steroidogenic enzyme gene *shade* in wings and, thus, controlling the synthesis of an essential growth hormone, 20E, directly in the wing tissue. In turn 20E signaling upregulates *Antp* expression. Antp also directly regulates the expression of critical cuticular protein genes in both forewings and hindwings.

**Fig 6.**
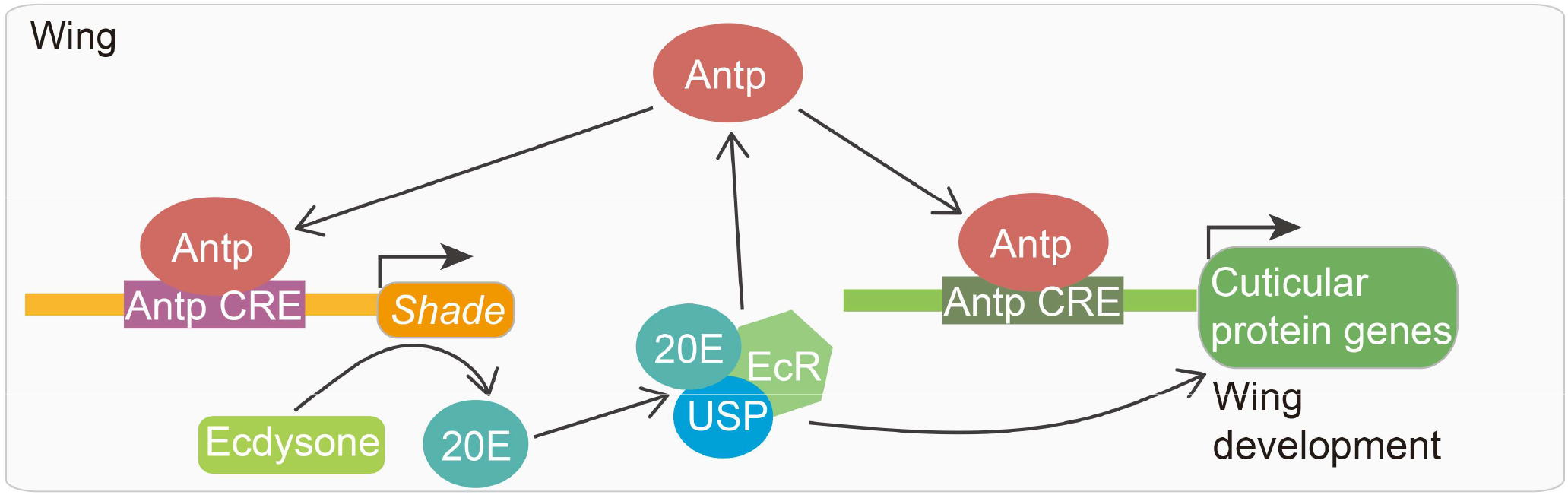
Proposed model on how *Antp* regulates wing development in *B. mori*. The Hox gene *Antp* plays an essential role in wing development. It does this by directly enhancing transcription of the steroidogenic enzyme gene *shade* in wings and, thus, controlling the synthesis of an essential growth hormone, 20E, directly in the wing tissue. In turn 20E signaling upregulates *Antp* expression. Antp also directly regulates the expression of critical cuticular protein genes in both forewings and hindwings.

## Materials and Methods

### Animal Strains

(1) The wild-type (WT) strain *DaZao* and mutant strain *Wes* (*Antp*^+/−^) were obtained from the Silkworm Gene Bank of Southwest University, China. Silkworms were reared on mulberry leaves at 25°C in ~75% relative humidity with a 12:12 h (L:D) photoperiod during their entire life. (2) The following fly stocks were used in this study: The WT *yw* and *nub-gal4* enhancer trap lines (BCF391#) were obtained from Core Facility of Drosophila Resource and Technology. The *UAS-Antp^RNAi^* (THU2760) was supplied by the Tisng Hua Fly Center. The wildtype *yw* were used as control flies. All individuals were incubated at 25°C. (3) The *Tribolium castaneum GA-1* strain was used in this study. Insects were reared in whole wheat flour containing 5% brewer’s yeast at 30°C under standard conditions.

### *Bombyx* cell lines

The *Bombyx mori* ovary-derived cell line BmN was cultured at 27°C in TC-100 medium (United States Biological) supplemented with 10% fetal bovine serum (Gibco) and 2% penicillin/streptomycin (Gibco).

### RNA Extraction and qRT-PCR

Total RNA samples were isolated from wing discs, prothoracic glands, hemolymph, BmN cells, and the whole beetles at different time points or under different conditions, using the MicroElute Total RNA kit (Omega) in accordance with manufacturer instructions. The cDNA was synthesized with 1 ug total RNA using the PrimeScript^TM^ RT Reagent Kit with gDNA Eraser (TaKaRa). qRT-PCR was performed using a qTOWER^3^G system (analytikjena) and a qPCR SYBR Green Master Mix (Yeasen). The eukaryotic translation initiation factor 4A (BmMDB probe ID sw22934) was used as an internal reference in *Bombyx*, and ribosomal protein S3 (*rps3*) in *Tribolium castaneum*. All experiments were independently performed with three biological replicates and the results were calculated using the 2-ΔΔCT method. Primers are listed in S1 Table.

### RNAi Experiment in *Bombyx* and *Tribolium*

The double-strand RNA (dsRNA) of *Antp, CPH28, ptl1, ptl2*, and *EGFP* were synthesized using the RiboMAX Large Scale RNA Production System T7 kit (Promega). Approximately 100 μg of synthesized ds*Antp* was injected into the second chest spiracle at the first day of *Bombyx* larval wandering stage. We injected 0.4–0.5 μg of *dsptl* at the ratio of 1:1 mix *ptl*1and *ptl*2 final instar larvae of *Tribolium castaneum*. To knockdown *CPH28* expression in the silkworm pupal stage, the siRNA sites 5’-GCAGCAAUUGUUCGCACAATT-3’ and 5’-GGAAGCUUUACAUUCGGUUTT-3’ (GenePharma) for CPH28 were designed. Ten μl of siRNA 1 μg/μl was injected from the breathing-valve into the wing disc on the 4^th^ day of the pupal stage. In addition, after injection, all insects were reared in a suitable living environment until analysis.

### Down-regulation of *Antp* in *Drosophila* wings

We used the Ga14-UAS system to knockdown *Antp* gene expression in *Drosophila* wings. We crossed the *UAS-Antp^RNAi^* males with *nub-gal4* virgin females and then incubated them at 25°C on a yeast/saccharose medium. The wing phenotypes of F1 adults were observed.

### CRISPR/Cas9-mediated *Antp* Knockout in *Bombyx*

The sgRNAs for knocking out *Antp* was designed by http://crispr.dbcls.jp/ and synthesized using the RiboMAXTM Large Scale RNA Production System T7 kit (Promega). Cas9 protein was purchased from Invitrogen (Thermo). The four sgRNAs and the Cas9 protein were mixed at a dose of 500 ng/ul. The mixture was incubated for 15 min at 37°C to produce a ribonuclearprotein complex (RNP) and micro-injected into the silkworm embryos within 2 h post oviposition. The injected embryos were incubated at 25°C and >90% relative humidity until they hatched. Genomic DNA of adult wings was extracted using the DNAzol (Takara) according to the manufacturer protocol. The target region was amplified using site-specific primers (Table S1). PCR products were checked by PAGE gel and sequencing approach. Related promoters are listed in Table S1. These sgRNAs synthesized *in vitro* were mixed with Cas9 protein and micro-injected into preblastoderm embryos of the *DaZao* strain.

### ELISA

ELISA was used to calibrate the ecdysteroid titer in wing disc of WT and Antp mutants. Silkworm wing discs were collected from ~50 pupae, and the pooled sample homogenized in methanol. The homogenate was centrifuged and we evaporated the supernatant at 55°C. The solid matter remaining was redissoved in 1 mL EIA buffer (Cayman Chemical) for 20E measurement and 1 mL sample diluents (BIOHJ) for ecdysone measurement, respectively. Ecdysteroid titers were assayed by an ELISA kit according to manufacturer instructions (Cayman Chemical or BIOHJ). Absorbance was measured at 414 nm for Cayman kit or 450 nm for BIOHJ kit on a BioTek H1 microplate reader.

### 20E Application

For 20E treatment in *Bombyx* and BmN cells, 20E (Adooq) was dissolved in DMSO and then diluted to the experimental concentrations with deionized distilled water. The final concentration of DMSO was 0.1% (v/v) in water. A total of 4 μg 20E was injected into larvae at the mesothoracic region on the 1st day of the larval wandering stage. An equal volume of DMSO at a final concentration of 0.1% (v/v) was used as the control. After 24 h, the wing discs were dissected in TRK lysis buffer (Omega). Five-μm 20E were applied to BmN cells for 24 h and then collected. An equal volume of DMSO was used as the control.

### Dual Luciferase Assay

The different lengths of *shade, CPH28*, and *Antp* promoters were subcloned into the pGL3-basic vector (Promega). The ORF of red fluorescent protein gene (RFP)-fused *Antp* was inserted into a pIZ/V5-His vector (Invitrogen) driven by the OpIE2 promoter. Different truncated promoters of pGL3-basic vector were co-transfected with pIZ/V5-His-Antp or treated with 20E at a concentration of 5 uM. After approximately 24 or 48 h transient transfection, dual-luciferase activities were measured using the Dual-Glo Luciferase Assay Kit (Promega). A pRL-TK vector containing the Renilla luciferase gene was used as an internal control.

### EMSA

Recombinant Antp nuclear proteins were extracted from *E. coli* strain BL21 (DE3) competent cells (TransGen). The potential Antp binding sites of the *shade* and *CPH28* promoters were predicted by the GENOMATIX system (http://www.genomatix.de/solutions/index.html) and JASPAR CORE (http://jaspar.genereg.net/). The DNA oligonucleotides containing Antp binding sites were labeled with biotin at the 5’-end and annealed to generate probes. EMSA experiments were conducted according to manufacturer instructions for the EMSA/Gel-Shift Kit (Beyotime). The binding reactions were performed with 4 μg recombinant Antp protein and different amounts of biotin-labeled probes (10 pmol, 20 pmol, 40 pmol) for 30 min at room temperature. For competition assays, 40 pmol unlabeled competitor probes were added to the reaction mixture. These samples were electrophoresed on 5% polyacrylamide gels in 0.5×TBE at room temperature. The total probes are listed in S1 Table.

### ChIP Assay

To further detect the effects of Antp on the activity of the *shade* and *CPH28* promoters, the ChIP assay was performed following kit instructions (GST). BmN cells were transfected with a *Flag-Antp* expression vector and harvested at 48 h. These cells were fixed with 37% formaldehyde, and then DNA containing proteins were sonicated to obtain 200–1000 bp length DNA fragments. The immunoprecipitation reactions were enriched with 1 μg antibody against Flag or IgG. The precipitated DNA and input were used for PCR analysis. The primers used for amplifying the sequences containing potential Antp binding sites are listed in S1 Table.

### Statistical Analysis

Statistical analyses were performed using GraphPad Prism 7 (GraphPad Software). The data are presented as the mean ± standard error (SE). The differences between two sets of data were analyzed with Student’s t-test. A value of P<0.05 was considered statistically significant; *P < 0.05, **P < 0.01, and ***P < 0.001.

## Acknowledgments

This work was supported by the National Natural Science Foundation of China [No. U20A2058, No. 31830094]. AM acknowledges support from the National Research Foundation Singapore Investigatorship award NRF-NRFI05-2019-0006.

## Author contributions

X. T. and C. F. designed the project. C. F., Y. X., and T. S. performed the experiment. C. F., A. M., and X. T. wrote the manuscript.

